# Recruitment of control and representational components of the semantic system during successful and unsuccessful access to complex factual knowledge

**DOI:** 10.1101/2021.06.02.446595

**Authors:** Silvia Ubaldi, Giuseppe Rabin, Scott L. Fairhall

## Abstract

Our ability to effectively retrieve complex semantic knowledge meaningfully impacts our daily lives, yet the neural processes that underly successful access and transient failures in access remain only partially understood. In this fMRI study, we contrast activation during successful semantic access, unsuccessful semantic access due to transient access-failures (i.e., ‘tip-of-the-tongue’, ‘feeling-of-knowing’), and trials where the semantic knowledge was not possessed. Twenty-four human participants (14 female) were presented 240 trivia-based questions relating to person, place, object or scholastic knowledge-domains. Analyses of the recall event indicated a relatively greater role of a dorsomedial section of the prefrontal cortex in unsuccessful semantic access and relatively greater recruitment of the pars orbitalis of the inferior frontal gyrus in successful access. Successful access was also associated with increased activation in knowledge-domain selective areas. Generally, knowledge-domain selective areas showed increased responses for both preferred and non-preferred stimulus classes. The exception was place-selective regions (PPA, TOS and RSC), which were recruited during unsuccessful access attempts for all stimulus domains. Collectively, these results suggest that prefrontal semantic control systems and classical spatial-knowledge selective regions work together to locate relevant information and that access to complex knowledge results in a broad activation of semantic representation extending to regions selective for other knowledge domains.

---

A defining characteristic of the human experience is our ability to learn and retrieve a broad range of complex factual knowledge. Our capacity to access this stored semantic knowledge impacts our ability to perform, scholastically, professionally and interpersonally, in our daily lives. Yet, the cortical processes underlying this capacity, and the mechanism underlying transient failures in our ability to access this knowledge, remain partially understood.

While we can find ourselves in a situation where we can successfully access a piece of knowledge, for instance, ‘what is the capital of Hungary?’, we can also find ourselves certain we know the answer but unable to retrieve it, only to be able to do so minutes, hours or days later. Unsuccessful semantic access (also called ‘blocking’; Schacter 1999) can be operationally characterised as a temporary failure in access, accompanied by accurate prediction of one’s capacity to recall the knowledge in a future test. Subjectively, this can be represented in the common experience of searching through our memory, navigating from related fact to related fact, in an attempt to get closer to the elusive piece of knowledge or lexical entry. Such states can be accompanied by an acute sensation that successful semantic access is imminent but just out of grasp, the so-called tip-of-the-tongue (ToT) state (Brown and McNeill 1966; Brown 1991), or the feeling that the knowledge is possessed without the accompanying sense of imminent retrieval (feeling-of-knowing; FoK; (Hart, 1967). Understanding these states can provide insight both into the neural substrates of these transient failure and into retrieval processes themselves.

Previous neuroimaging work has highlighted increased activation of bilateral prefrontal control systems when comparing unsuccessful access to a combination of successful and knowledge-absent trials (Maril et al. 2001, 2005) or when parametrically comparing unsuccessful semantic-access trials knowledge-absent trials (Kikyo et al. 2002). The identified prefrontal regions include those implicated in guiding the retrieval of semantic control (left inferior frontal gyrus (IFG), dorsomedial prefrontal cortex - supplementary motor area (dmPFC-SMA); Jackson 2021), as well as those that are not (right IFG, bilateral anterior PFC). One potential consideration is that studies comparing known to blocked states have included in the modelled fMRI response both the recall-event and the subsequent metacognitive evaluation of the participants internal cognitive state during the recall-event. Consequently, the involvement of frontal regions may reflect extra-semantic processes, such as response selection of cognitive conflict associated with the metacognitive judgement (Maril et al. 2005).

At the same time, the brain’s semantic system includes an extensive left-lateralised set of brain regions that reliably activate more strongly to semantically richer stimuli (Binder et al. 2009), including regions showing univariate and multivariate sensitivity to the semantic content of probe stimuli that may be more closely linked to the representation of semantic knowledge (Fairhall and Caramazza 2013a; Fernandino et al. 2015; Liuzzi et al. 2017, 2020, 2021). Failures in semantic access may also arise from a failure to ignite such semantic representations.

The objective of the current study is to gain insight both into the processes underlying temporary blocks in semantic and lexical access and those that allow successful access to stored knowledge. We contrast the fMRI response associated with successful and unsuccessful semantic access, and trials where semantic knowledge is absent. This study extends the existing literature in two ways. Firstly, we employ an fMRI design that permits the deconvolution of the cortical responses to the cognitive recall-event from subsequent metacognitive decisions about the degree of confidence that the knowledge is possessed. Secondly, we further examine the role of representational components of the semantic system in successful and unsuccessful semantic access. To accomplish this, we utilise general knowledge questions drawn from four different knowledge domains. This manipulation of the semantic content of the probe stimuli allowed us to better isolate regions involved in representational processes as well as to understand the relationship between knowledge-domain selectivity in the cortex and successful and unsuccessful semantic access.

## MATERIALS AND METHODS

### Participants

Twenty-nine right-handed native-Italian speakers with no history of neurological disorder were recruited for the study. Three participants were excluded due to within-run head motion exceeding 2 mm or excessive ocular artifacts during scanning (excessive condition-specific signal emanating from the eyes). One subject was excluded as they did not report any of the place or scholastic facts to be unknown and a further participant did not complete the entire experiment. Thus, the final sample included 24 participants (14 female, mean age 26.4 years). Participants gave informed consent and were compensated for participation (15 €/hour). The study was conducted in line with the declaration of Helsinki (1964, amended in 2013) and was approved by the Ethics Committee at the University of Trento.

### Experimental Design

#### Stimuli

Stimuli were composed of 240 Italian written general-knowledge questions. The questions were equally divided into four knowledge domains: People, in which there were questions about famous historic or fantasy characters or famous contemporary people (e.g. ‘*Which philosopher uttered the phrase I know I don’t know?’*); Places, where they were asked about famous geographic places or monuments (e.g. *‘In which Spanish city is the Alhambra complex located?’*); Objects, in which the questions were about specific objects or materials (e.g. *‘What is the name of the element that supports the weight of an arch?’*). The final domain of knowledge, ‘scholastic’, was designed to capture category-general factual knowledge that was unrelated to direct experience with the environment, more likely to be learned verbally, and did not involve person-, place or object-related information. (e.g. *‘What is the name of the transition of matter from the solid to the gaseous state?’*).

Stimuli were matched across knowledge-domain by number of words (F_(3,236)_=1.13, p=.337) and number of letters (F_(3,236)_=1.12, p=.343). The full list of stimuli is available here: https://figshare.com/s/194aa97ca67800f4c658.

#### fMRI Experimental Task

The fMRI session was divided into 4 experimental runs. In each run, fifteen questions from each knowledge-domain were presented in a pseudo-randomized order using Matlab (www.mathworks.com) and Psychtoolbox Version 3 (www.psychtoolbox.org). Each trial was divided into two parts. Firstly, the recall-event included the 3-second written presentation of the question followed by a 3-second fixation cross. Participants were instructed to read the question once and indicated, via two-option button press with their right hand, whether the knowledge was accessible or inaccessible (non-responses (<1%) were coded as inaccessible). If participants indicated they could access the answer, the experiment continued to the next trial. Inaccessible response prompted a 0 to 8 seconds jitter followed by a metacognitive judgement. Here, participants were cued to indicate, via three-option button press with the same hand, whether they experienced a TOT, a FOK, or that they answer was unknown (4 second duration). Participants were instructed to respond ‘known’ if they could access the answer in that moment, to indicate a ToT if they were convinced they knew the answer and were on the verge of producing it but could not quite do so, to respond FoK when, while the answer was not presently accessible, they felt they knew it and were certain they would be able recognise the correct answer, and to respond ‘unknown’ if they did not think the they knew the answer.

#### Post Scanner Knowledge-Verification Test

To validate the self-reported responses in the scanner, after the fMRI session participants were again presented with the questions classified as ToT, FoK or unknown, as well as 20% of the ‘known’ questions. Participants were allowed 5 seconds to type the answer to each question without any cue. If the participant did not respond or the participant opted to proceed to the cue (via button press), the first letter of the answer appeared, and they had an additional 5 seconds to start typing the response.

### MRI scanning parameters

Functional and structural data were collected at the Center for Mind/Brain Sciences (CIMeC), University of Trento, with a Prisma 3T scanner (Siemens AG, Erlangen, Germany), using a 64-channel head coil. Participants lay in the scanner and viewed the visual stimuli through a mirror system connected to a 42”, MR-compatible, Nordic NeuroLab LCD monitor positioned at the back of the magnet bore. Functional images were acquired using echo planar (EPI) T2*-weighted scans. Acquisition parameters were: repetition time (TR) of 2 s, an echo time (TE) of 28 ms, a flip angle of 75°, a field of view (FoV) of 100 mm, and a matrix size of 100 × 100. Each volume consisted of 78 axial slices (which covered the whole brain) with a thickness of 2 mm, AC/PC aligned. High-resolution (1×1×1 mm) T1-weighted MPRAGE sequences were also collected (sagittal slice orientation, centric phase encoding, image matrix = 288 × 288, field of view = 288 mm, 208 slices with 1-mm thickness, repetition time = 2290, echo time = 2.74, TI = 950 ms, 12° flip angle).

### fMRI Data Analysis

Data were analysed and preprocessed with SPM12 (http://www.fil.ion.ucl.ac.uk/spm/). The first 4 volumes of each run were dummy scans. All images were slice-time corrected, realigned to correct for head movement, normalised to MNI space and smoothed using a 6mm FWHM isotropic kernel. Before computing the general linear model (GLM), the four runs were concatenated, to avoid empty parameters in one or more conditions. Sixteen trial types were modelled for the recall event: the four different recall-types (successful-access, ToT, FoK and knowledge-absent trials) for each of the four knowledge domains. The type of these different recall-events, for ToT, FoK and knowledge-absent trials, was determined by re-coding based on the subsequent metacognitive judgment. Twelve additional regressors were included to capture the metacognitive part of the trials, with TOT, FOK and knowledge-absent responses modelled for each of the four categories. Subject-specific parameter estimates (β weights) for each of these 28 conditions were derived through the GLM. The 6 head-motion parameters were included as additional regressors of no interest. Group level analysis was performed in two separate random effects GLMs, one for the recall event and one for the metacognitive judgment.

Group level whole-brain analyses of response type was performed using weighted contrasts for: unsuccessful access [(ToT&FoK)>knowedge-absent] and successful access [known>(ToT&FoK)]. Domain-selective responses were taken by contrasting each domain with the average of the other three domains. Analysis of the metacognitive judgement was determined by contrasting ToT, FoK and unknown responses with one another. All analyses were performed within an inclusive mask designed to isolate regions previously implicated in semantic processing (see results). Unless otherwise specified, results are reported at an initial voxel-wise threshold of p<.001, with family wise error (FWE) correction for multiple comparisons at the cluster level (p<.05) as implemented in SPM12.

Two region-of-interest (ROI) analyses were performed to assess: 1) the effect of recall type in semantic control systems and, 2) the effect of recall type within knowledge-domain selective brain regions. In the former, 5mm spherical ROIs were constructed in the pars orbitalis and pars triangularis sections of the IFG and in the dmPFC-SMA based on coordinates previous published in a recent meta-analysis of semantic control processes (Jackson, 2021). In the latter, ROIs were defined via the domain-selective contrasts (e.g. place > mean(person, object, scholastic)). As the ROI-defining contrasts contain no information about response type, subsequent analyses of response type or response-type by domain interactions are orthogonal and statistically unbiased (Friston et al. 2006) These knowledge-domains selective ROIs were constructed by taking the union between a sphere with a radius of 5mm centred at the selectivity peak for the knowledge-domain (see table 2) and statistically significant voxels for the relevant domain-selective contrast (see figure 4). Within knowledge-domain selective ROIs, data were analysed for all domains, preferred domain, or the average of the non-preferred domains in a series of weighted contrasts that isolated the effects of interest, as detailed in the relevant section of the results.

## Results

### Behavioural

Participants reported that the fact was known on 45.6% of trials, ToT states on 12.7% of trials, FoK states on 16.3%, and that the fact was unknown on 25.5%, in keeping with previous results using similar paradigms (Maril et al. 2001, 2005). Post-scanner testing indicated that known responses were genuine, with 79.5% of correct responses being correctly provided, either through free recall (48.8%) or when cued by the first letter of the word (30.8%). Accuracy for ToTs (60.0%; 32.3% unprompted, 27.2% cued) were significantly lower (t_(23)_=5.33,p <.0001), and accuracy was for ToT responses (38.0%; 16.9% unprompted, 21.1% cued) were progressively lower (t_(23)_=7.41,p <.0001). Finally, subjects could accurately report 20.4% of unknown responses (8% unprompted, 11.9% cued; significantly lower than FoKs, t_(23)_=6.56,p <.0001).

To assess RTs for the initial recall event (measured from the onset of the probe sentence), the responses were decomposed based on the subsequent metacognitive judgment. The effects of response type (known, ToT, FoK and knowledge-absent) and knowledge domain (people, places, objects and scholastic) were then assessed via repeated measures ANOVA. RTs differed as a function of response type (F_(3,69)_=41.6, p<.001). Paired t-test revealed that ToT (3797 msec) and FoK (3730 msec) did not significantly differ (t_(23)_=1.57, p=.13) and were slower than responses subsequently confirmed to be unknown (3464 msec; v. ToT t_(23)_=5.43, p<.001; v. FoK t_(23)_=5.08, p<.001). Unknown responses were in turn slower than known responses (3238 msec; t_(23)_=4.97, p<.001). RTs also differed as a function of knowledge domain (F_(3,69)_=28.3, p<.001). Judgments for people trials (3361 msec) were faster than for places (3548 msec; t_(23)_=4.31, p<.001), which were in turn faster than objects (3670 msec; t_(23)_=3.49, p=.002) and scholastic items (3651 msec, t_(23)_=2.84, p=.009), which did not differ significantly from one another (t_(23)_=0.64, p=.53). There was no interaction between response type and knowledge domain (F_(9,207)_=.99, p=.45).

RTs for the metacognitive judgment differed modestly as a function of category (F_(3,69)_=2.88, p=.042), with faster responses for people (559 msec) than for object (618 msec, t_(23)_=2.42, p=.024) and scholastic (604 msec, t_(23)_=2.46, p=.022) domains. Response type had a more pronounced influence on RTs (F_(2,46)_=10.63, p<.001), with slower responses for FoK (673 msec) compared to both ToT (569 msec, t_(23)_=3.59, p=.002) and unknown (560 msec, t_(23)_=5.90, p=.001) responses.

### Inclusive Masking

Unexpectedly, when the answer to a question was thought to be known ([successful or unsuccessful access] > knowledge-absent), strong activation was seen in regions associated with reward, bilateral caudate, anterior insula and the substantia nigra of the midbrain (see also, Kikyo et al. 2002). As the contribution of these regions is uncertain, and to focus analyses on regions previously implicated in semantic processing, subsequent analyses have been restricted to regions showing sensitivity to semantic category at an uncorrected (p<.05) threshold (omnibus ANOVA: main effect of category, see figure 1). At this lenient threshold, the mask can be seen to encompass both control and representational aspects of the semantic system. Examination of excluded regions showing a stronger response on trials where the response was thought to be known with Neurosynth image decoder (https://neurosynth.org/decode/; Yarkoni et al. 2011; Rubin et al. 2017), indicated correspondence with reward-related terms (gain=.359, incentive=0.263, reward=0.252; three highest non-anatomic reverse-inference term). Full details of the excluded regions are available here: https://figshare.com/s/2f4be0ba0278ea79a7d5.

**Figure 1.**
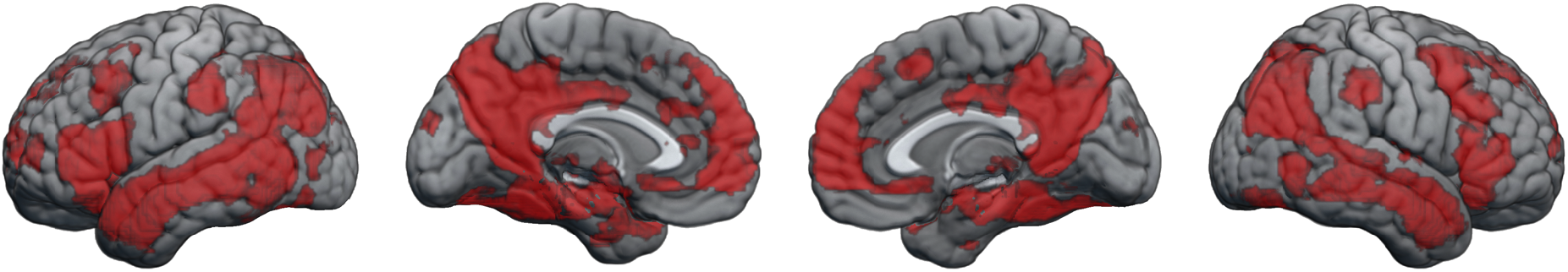
Inclusive functional mask applied to all contrasts in this study. The mask was defined as regions showing sensitivity to knowledge-domain at a lenient statistical threshold (p<05, uncorrected).

### Neural correlated of blocked and successful semantic access

Within the inclusive mask, we first compared unsuccessful retrieval (ToT/FoK) to trials where the knowledge was absent (figure 2 and table 1). Relative to knowledge-absent trials, unsuccessful access recruited semantic control circuitry in the left lateral prefrontal cortex and medial supplementary motor cortex. Additionally, the bilateral retrosplenial complex (RSC), left parahippocampal gyrus, the transverse occipital sulcus, and a region of the supramarginal gyrus, were more active during FoK/ToT than knowledge-absent trials.

**Figure 2.**
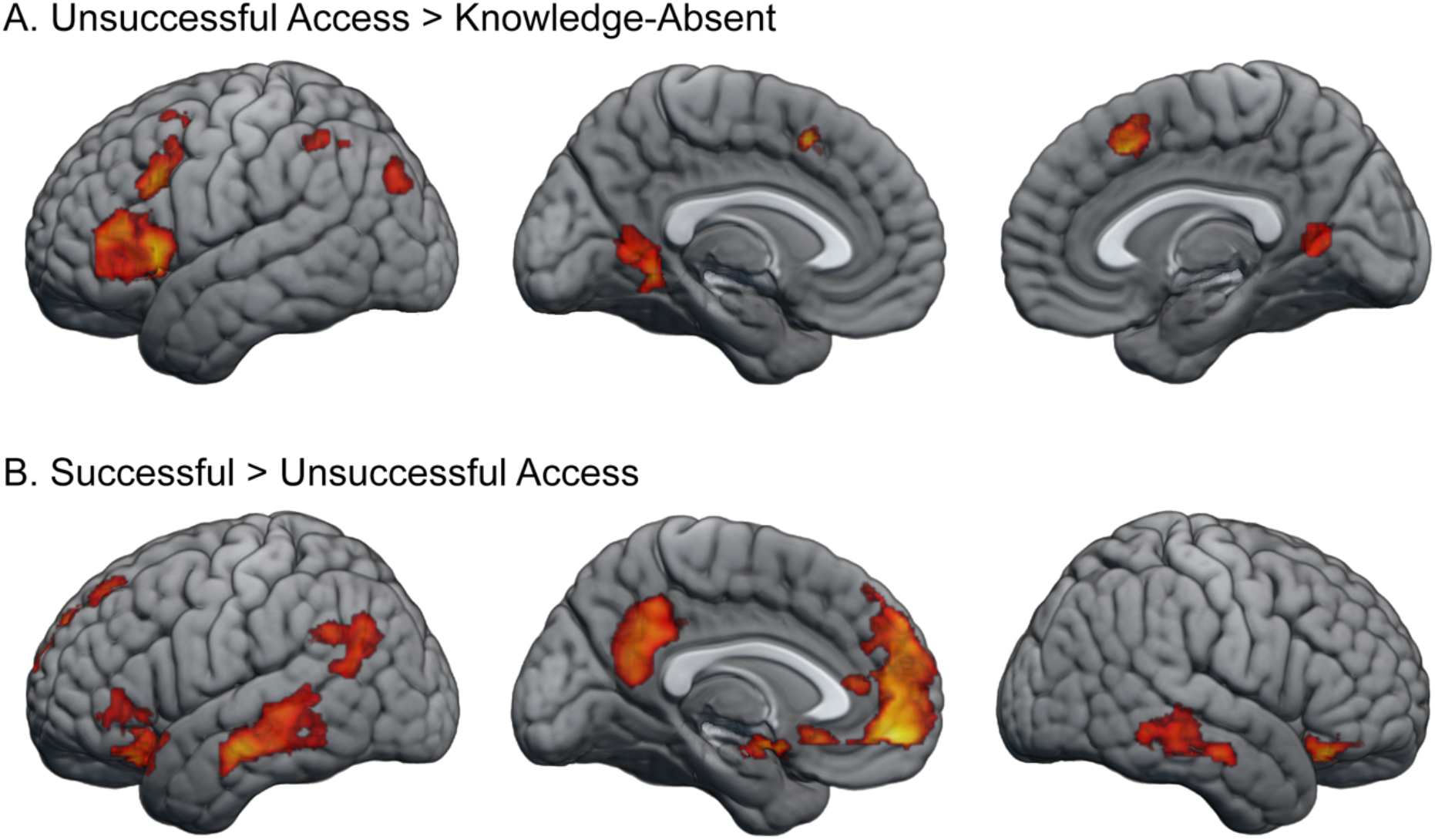
Successful and unsuccessful access during the recall-event. Shown is the response to A) unsuccessful semantic access compared to trials where the knowledge was absent (ToT and FoK trials > knowledge-absent trials) and B) Successful > Unsuccessful access. Brain maps show the response during the recall event, when participants indicated whether the knowledge was accessible or inaccessible (with inaccessible responses coded into ToT, FoK or knowledge-absent trials based on the subsequent metacognitive judgement). Brain maps were thresholded with an initial voxel-wise threshold of p<.001 and cluster corrected at p<.05. See table 1 for significance, extent and location of clusters.

**Table 1.**
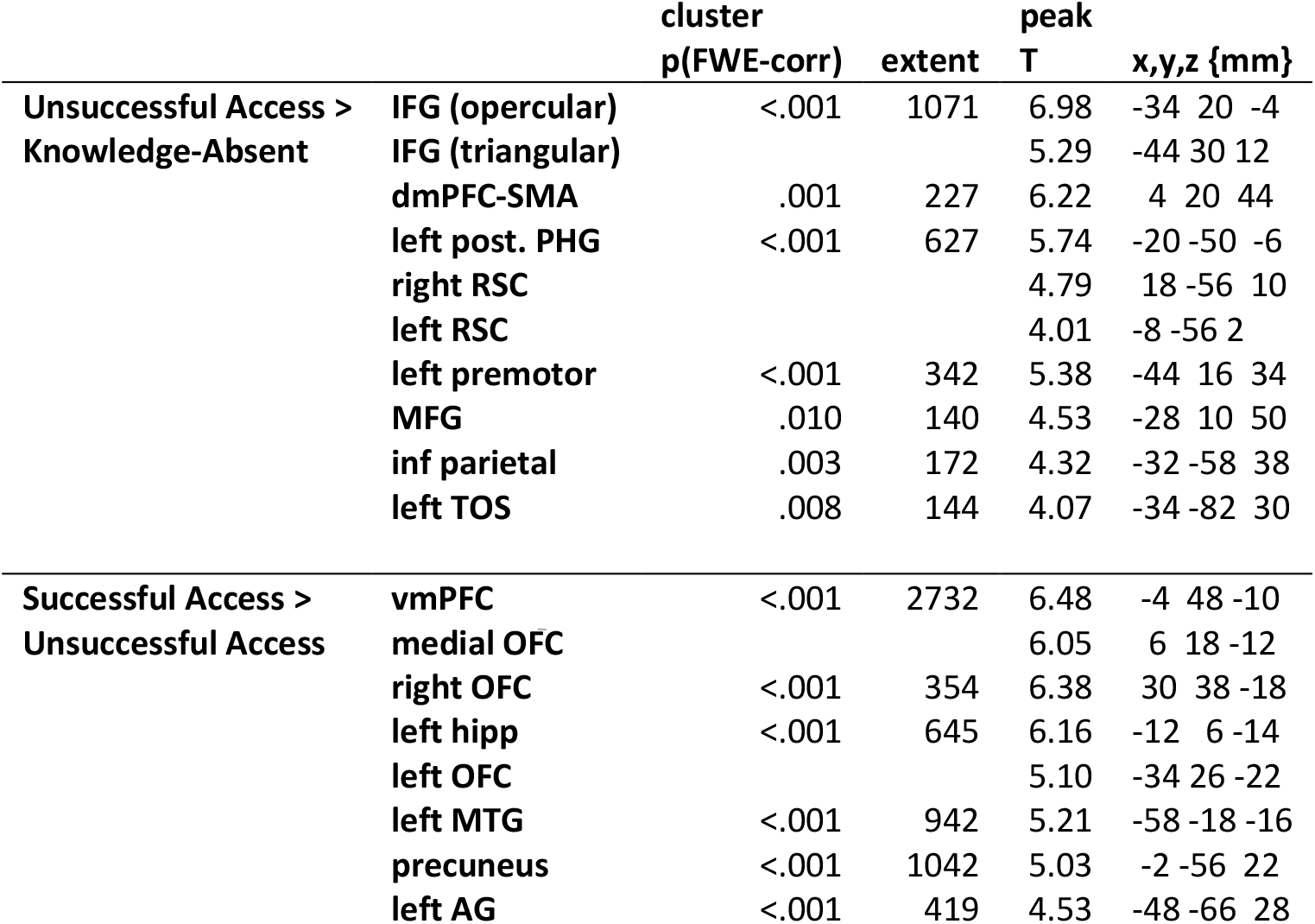
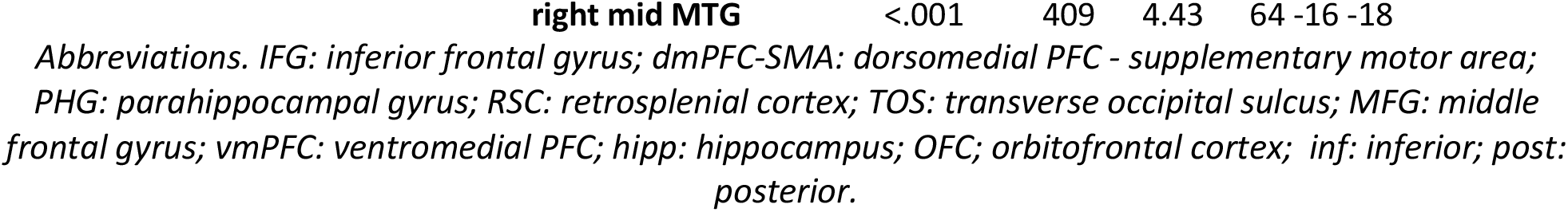
Significance, extent and location of regions showing differential responses for successful and unsuccessful semantic access as shown in figure 2. For large clusters spanning multiple regions, local maxima are listed separately.

**Table 2.**
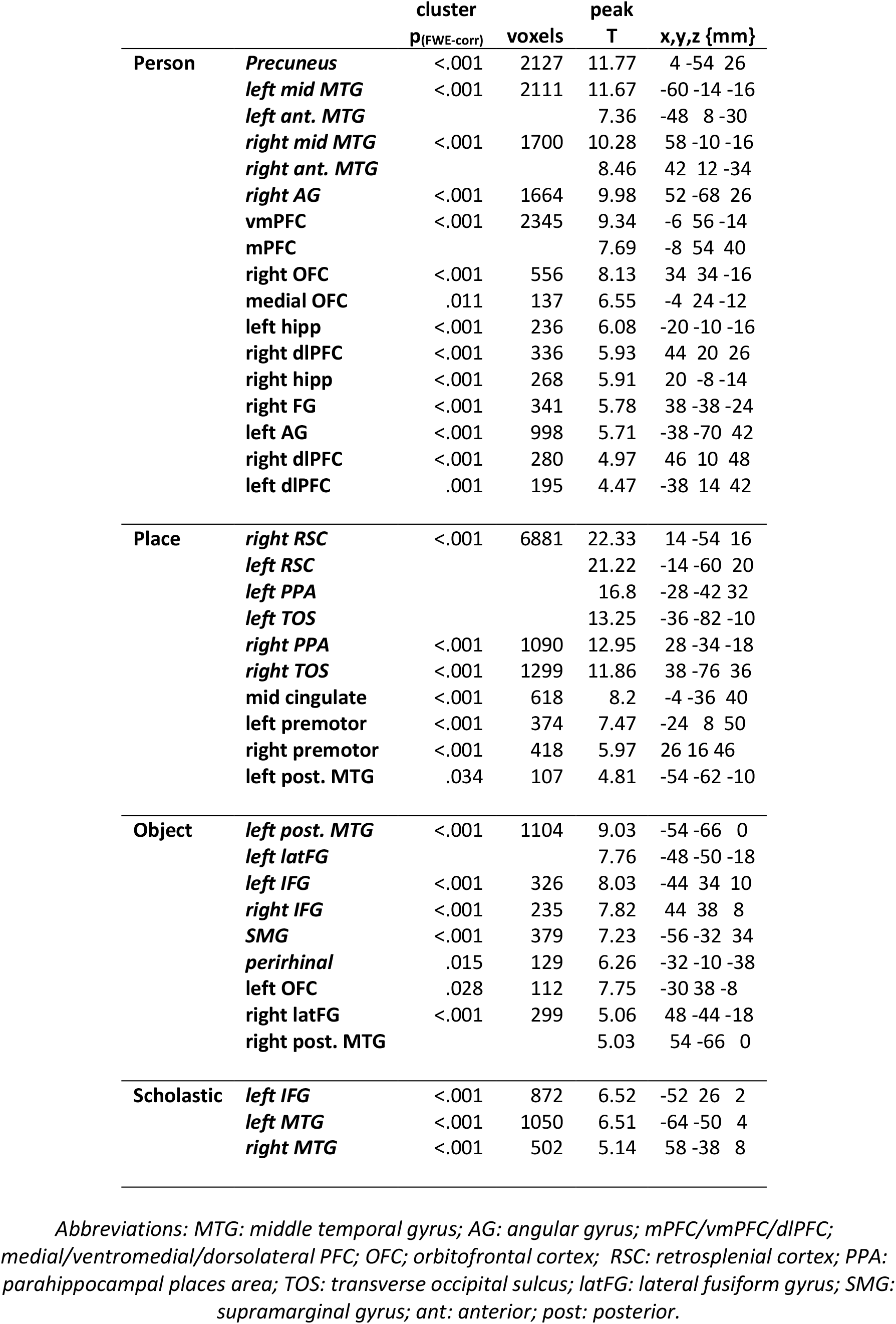
Significance, extent and location of regions showing differential responses for knowledge domain, as shown in figure 4. For large clusters spanning multiple regions, local maxima are listed separately. Italicised regions are incorporated in subsequent ROI analyses.

To delineate the neural processes associated with successful semantic access from failed semantic access, we contrasted successful to unsuccessful (ToT/FoK) access. Results show increased cortical activity in a network of regions associated with the representation of semantic knowledge: the precuneus, mPFC, bilateral sections of the middle temporal gyrus and the left angular gyrus and hippocampus. Additionally, lateral orbitofrontal cortex in both hemispheres were seen to be more active when stored information was successfully accessed. Notably, activation in the left OFC was seen to extended laterally into the pars triangularis of the IFG.

No regions showed a significant difference between ToT and FoK trials during the recall event. Trials counts were relatively low for ToT and FoK trials, which may reduce statistical power and it is possible that true differences between these response types exist during the recall event that were below the ability of present paradigm to resolve (but see later section: metacognitive judgements of knowledge confidence).

Reaction times were shorter in successful-access that unsuccessful-access trials and increased cortical responses for successful access are unlikely to be attributable to increased RT or effort. However, increased responses for unsuccessful compared to knowledge-absent trials may be attributable to the longer RTs associated with the former. In a supplementary analysis, we considered this issue by including the RT of each trial as a covariate at the first level GLM. At the a priori statistical threshold (p<.001 voxel, p<05, cluster corrected), differences between unsuccessful and knowledge-absent trials were not evident. Due to the systematic variations in RT between conditions, the covariate is expected to dilute statistical differences, and this is expected even if RT is not driving cortical effects. At an exploratory threshold of p<.005 voxel level with the same inferential threshold (p<.05, cluster corrected), effects were present in all reported clusters (p-values <.008, cluster-corrected), with the exception of the dmPFC-SMA (p=.148). In this way, we can be reasonably confident that the reported difference between unsuccessful and knowledge-absent trials in those regions are not solely attributable to RT differences.

To more fully characterise the differences between response types in prefrontal semantic control regions, unbiased ROI analysis was performed based on published coordinates. Specifically, 5mm radius spheres were constructed around peak coordinates for the left IFG pars triangularis (MNI=-48, 22, 20), pars orbitalis (MNI=-46, 24, -2) and the dmPFC-SMA (MNI=-2 20 52), as reported in a recent meta-analysis of semantic control studies (Jackson, 2021). As expected from the whole-brain analysis, the response was greater for successful access and unsuccessful access than knowledge-absent trials in all regions (figure 3). However, different profiles were evident across the three regions (region by response type interaction: F_(4,92)_=12.16, p<.0001) and when each region was compared to each other (IFG_orb_<-> IFG_tri_: F_(2,46)_=6.53, p=.003; IFG_tri_ <–> dmPFC-SMA; F_(2,46)_= 7.49, p=.002; IFG_orb_ <–> dmPFC-SMA (F_(2,46)_=21.32, p<.0001). While responses were equivalent for successful and unsuccessful access in the left IFG pars triangularis (t<1), responses in the dmPFC-SMA were stronger during unsuccessful compared to successful access (t_(23)_=3.36, p=.003) and stronger for successful than unsuccessful access in the pars orbitalis (t_(23)_=2.94, p=.007). The dissociation between regions persisted when comparing only successful and unsuccessful trials (IFG_orb_ <-> IFG_tri_: F_(1,23)_=9.46 p=.005; IFG_tri_ <–> dmPFC-SMA; F_(1,23)_= 13.81, p=.001; IFG_orb_ <–> dmPFC-SMA (F_(1,23)_=35.33, p<.0001). Collectively, these results indicate robust differences between semantic control regions in their involvement in successful and unsuccessful semantic access. Follow-up analysis of differences between ToT and FoK trials revealed no significant differences in the left IFG pars triangularis (t<1), left IFG orbitalis (t_(23)_=1.17, p=.25) or dmPFC-SMA (t_(23)_=1.72, p=.10).

**Figure 3.**
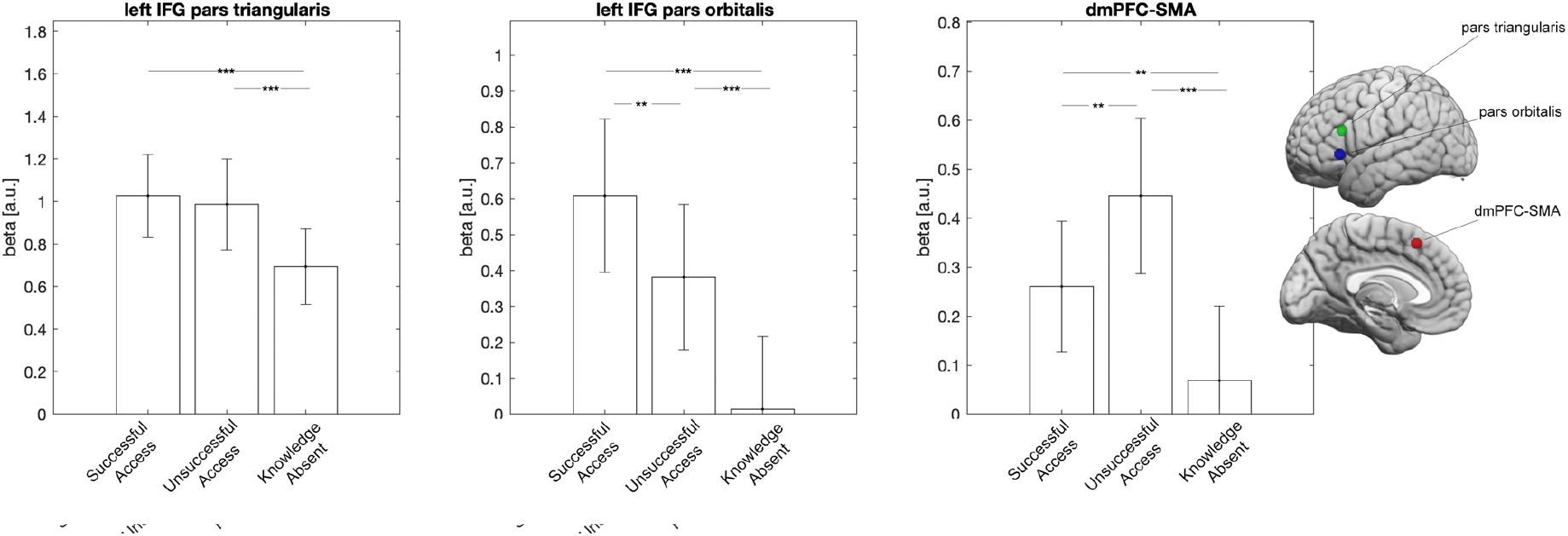
Region-of-interest analysis of independently localised semantical-control regions. Shown are the fMRI response magnitudes as a function of response type for the left IFG pars triangularis (green), pars orbitalis (blue) and the dmPFC-SMA (red). Error bars indicate ±1 between-subject standard error of the mean. Asterixis indicate within-subject statistical significance (***<.001, **<.01, *<.05).

**Figure 4.**
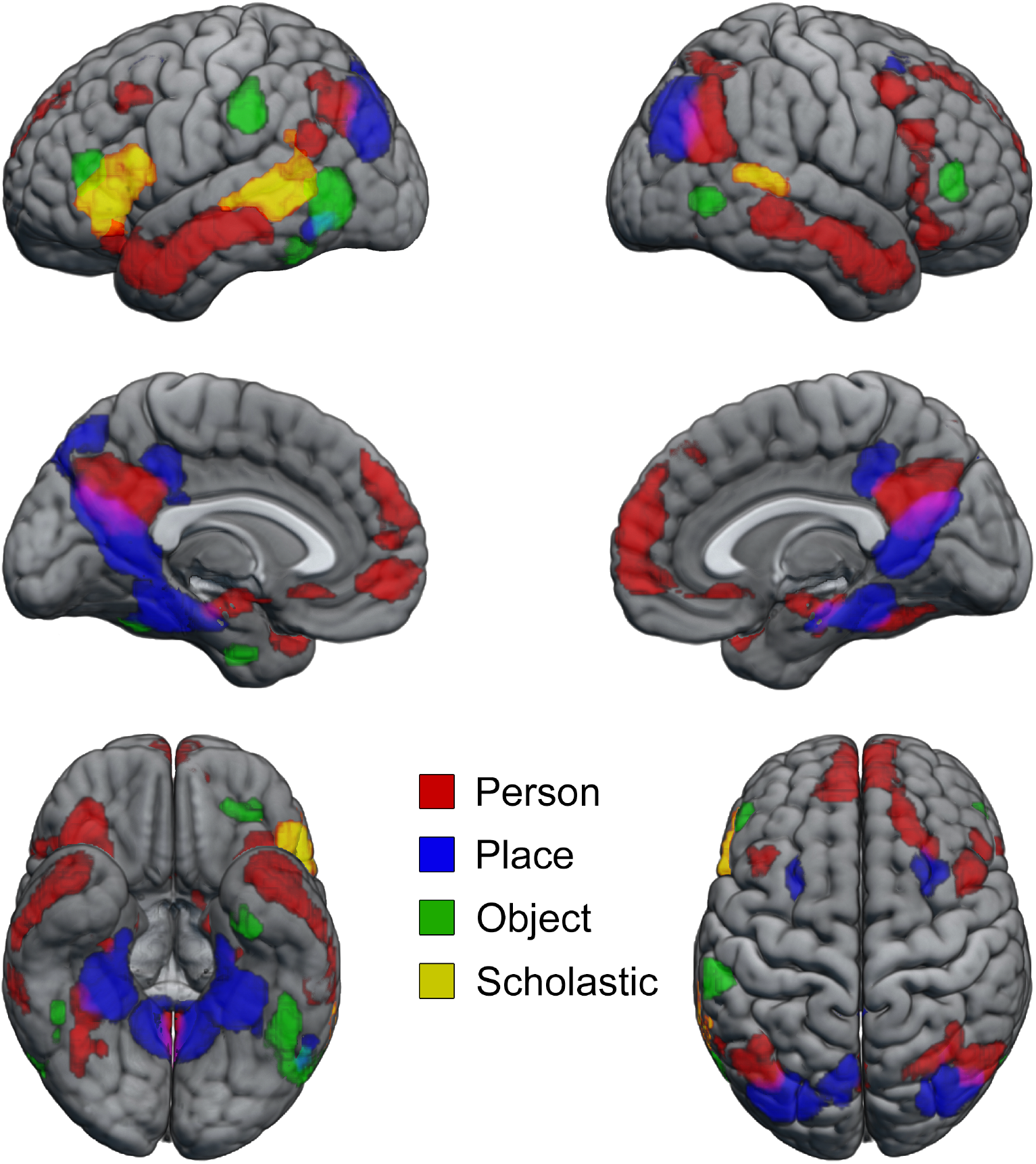
Cortical regions demonstrating a preference for one category versus the average of the others during the recall attempt, irrespective of recall outcome. Brain maps are initially thresholded at a voxel-wise threshold of p<.001 and cluster-level corrected p_FWE_<.05. See table 2 for significance, extent and location of clusters.

### Comparing unsuccessful to successful semantic access

Whole brain analysis of the contrast of unsuccessful > successful access identified two small clusters. One in the left premotor area in a region overlapping with the frontal eye-fields (extent: 174 voxels; p_corr_=.003; *xyz*_MNI_: -22 -2 52) and another in the left superior parietal lobe (extent: 163 voxels; p_corr_=.004; *xyz*_MNI_: -8 -72 52). Follow up investigation showed that these regions were not more responsive for unsuccessful access compared to knowledge-absent trials and the nature of this effect remains ambiguous.

### Knowledge-domain selective responses

To identify brain regions sensitive to semantic content, each knowledge domain was contrasted to the average of the other three knowledge domains. Knowledge-domain selectivity was evident in an extensive set of temporal, parietal and prefrontal regions (Figure 4, Table 2). Person knowledge selectively activated bilaterally: mid and anterior middle temporal gyrus, angular gyrus, premotor cortex and hippocampi; medially: the precuneus and mPFC; as well as the lateral fusiform gyrus in the right hemisphere. In addition to the classic place-selective network, (PPA, RSC, TOS) place stimuli more strongly recruited superior aspects of the prefrontal cortex bilaterally, left pMTG and the midcingulate cortex. Notably, three regions outside of the semantic control network which show an increased response during blocked semantic access (figure 2A), correspond to left PPA and TOS and bilateral RSC.

Object-related retrieval events selectively activated bilateral IFG, pMTG and lateral fusiform gyri as well as the left supramarginal gyrus and perirhinal cortex. Scholastic retrieval events selectively activated left IFG and the posterior superior temporal sulcus bilaterally. The degree of overlap in voxels showing a preference for more than one domain (indicated by blended colourmaps in figure 4) was minimal. Notably, reaction times of the conditions comprising the domain-selective contrasts associated with the longest RTs, object and scholastic, did not significantly differ (t<1). Consequently, increased RTs alone are unlikely to account for domain-selective seen between these domains.

### Effect of recall-type within domain-selective brain regions

To investigate the influence of recall type within brain regions exhibiting a sensitivity to semantic content, ROIs were defined via the orthogonal domain-selective contrasts presented in figure 4 and table 2. To reduce multiple comparisons over the large number of identified domain-selective regions, analyses were performed by averaging the responses of ROIs within each network. Specifically, the six most strongly selective regions (indicated in italics in table 2) were selected for person, place and object domains and all three regions for the scholastic domain. This approach was adopted to maximise signal and balance the number of voxels contributing to each network.

Within person, object and scholastic domain-selective networks, an overall increase in activity irrespective of sentence domain was seen from knowledge-absent to successful-access trials (all p-values<.0001) and from unsuccessful access to successful access trials (all p-values<.005; see figure 5A). Increased activation during unsuccessful access compared to knowledge-absent trials was largely absent, with modest effects limited to the object-domain-selective network (p=.031). Notably, within these domain-selective networks, the modulation by recall-type was not sensitive to stimulus domain. For comparison, figure 5B shows the effect of recall-type for each network’s preferred stimulus type and figure 5C shows the response for the average of the three non-preferred stimulus classes. Descriptively, the response profile across different recall-types in person, object and scholastic domain-selective networks can be seen to be largely similar for preferred and non-preferred stimulus classes (compare figure 5B & 5C). Inferentially, this was assessed through weighted contrasts which demonstrated that increased responses during successful compared to unsuccessful access were in fact larger for non-preferred compered to preferred domains in person (t_(23)_=6.38, p<.001), object (t_(23)_=3.63, p=.001), and scholastic (t_(23)_=2.21, p=.045) networks. This indicates that, in person, object and scholastic domain-knowledge-selective regions, the enhanced response associated with successful access was not domain-selective.

**Figure 5.**
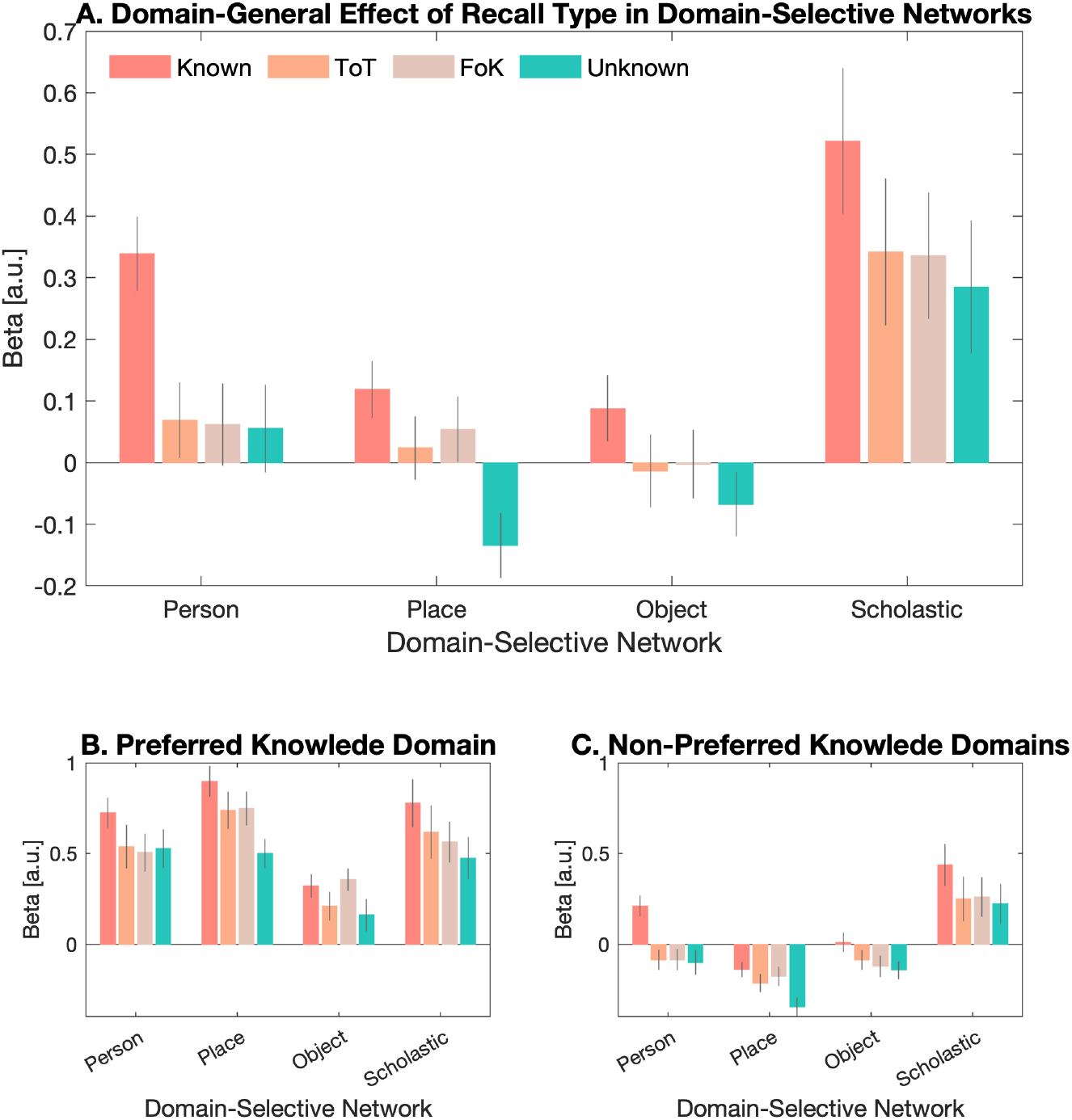
Effect of Recall Type within Knowledge-Domain Selective Cortical Networks. **A**. Effect of Recall Type for all stimulus domains. **B**. Effect of Recall Type for each network’s preferred domain (e.g., person-related questions in person-selective regions). **C**. Effect of Recall type for each network’s non-preferred domain (e.g., the average of place-, object-, scholastic-related questions in person-selective regions). Error bars indicate ±1 standard error of the mean.

A different profile was evident within the place-selective network. Like the other domain-selective networks, a domain-general increase was present from unknown to known trials (p<.0001). Unlike the other networks, the increased response from knowledge-absent to successful-access trials was domain selective. Specifically, there was a greater increase for place-related trials compared to the other domains (p=.002) and there was no significant increase from unsuccessful-access to successful-access trials when only the three non-preferred categories were considered (p=.1). In addition, place-selective regions exhibited a robust increase in response for unsuccessful-access trials relative to knowledge-absent trials (p=.0002), an effect that persisted when only non-preferred stimuli were considered (p=.004). To assess the specificity of increased activity during unsuccessful compared to knowledge-absent access trials to the place selective network, repeated measured ANOVA was performed comparing the strength of the effect (unsuccessful minus knowledge-absent trials) across the four networks. A significant effect of region (F_(3,69)_=5.24, p=.003) was driven by significantly stronger effects in the place-selective network than people (T_(23)_=5.32, p<.0001), objects (T_(23)_=3.09, p=.005) selective networks, and stronger effects that approached in scholastic knowledge-domain selective network (T_(23)_=2.07, p=.050), This results confirms the greater involvement of the place-selective network in unsuccessful access.

While averaging across ROIs within a given network reduces multiple comparisons, it introduces the risk of mistaking local regional effects for global network-level effects. In a supplementary analysis, differences within the ROIs composing the knowledge-domain selective networks were assessed via repeated measures ANOVA, following weighted averaging of the conditions to isolate the difference between successful versus unsuccessful access and between unsuccessful access versus knowledge-absent trials.

Comparing successful to unsuccessful access, interregional differences were limited to the person-domain network (f_(5,115)_=6.01, p<.001). Post hoc paired t-tests revealed that this was driven by larger effects in vmPFC compared to other regions, with differences between vmPFC and left lateral ATL, rTPJ and left mid-MTG, surviving Bonferroni correction for multiple comparisons across regions (15 comparisons: adjusted alpha = .0033). However, each region individually showed greater responses for successful than unsuccessful access (p-values <.0001, adjusted alpha = .0083, indicating that, despite that variations across the network, all regions are contributing to the reported network-level effects. Comparing unsuccessful access to knowledge-absent trials, interregional differences in the object selective (F_(5,115)_=11.26, p<.001) and scholastic (F_(2,46)_=16.50, p<.001) networks were both driven by stronger response in the left IFG compared to other regions (object network: p-values<.001, α=.0083; scholastic network: p-values< .001, α=.0167). This pattern is consistent with the whole brain analysis and role of this region in semantic control (figure 2). Finally, differences in the place-domain network (F_(5,115)_=4.13, p=.0017) were driven by non-significant increases for unsuccessful access trial in right PPA, that were significant smaller than effects in left PPA and right TOS (p-values<.0025, α=.0083), suggesting the right PPA is less involved than other elements of this network in attempts to access semantic knowledge.”.

### Metacognitive judgments of knowledge confidence

To examine the neural correlates associated with the metacognitive judgement of the degree of confidence with which inaccessible knowledge was believed to be possessed, we compared ToTs and FoKs to unknown responses in the metacognitive stage of the trial. Both ToTs and FoKs activated prefrontal control regions bilaterally and medially (see figure 6). There was a graduated response, with greater responses for ToTs than FoKs. Specifically, all regions showing a response to FoKs also exhibited a response to ToTs, activation of control circuitry was more expansive for ToTs than FoKs and was significantly stronger in bilateral IFG and the right middle frontal gyrus, indicating that it was possible to distinguish ToT from FoK states within the current paradigm. ToTs additionally activated representational components of the semantic system the lateral mid-MTG bilaterally and the left temporoparietal junction. Despite longer reaction times for FoKs that ToTs, no region responded more strongly to FoKs than ToTs, indicating that these effects are not driven by reaction time differences or general level of difficulty

**Figure 6.**
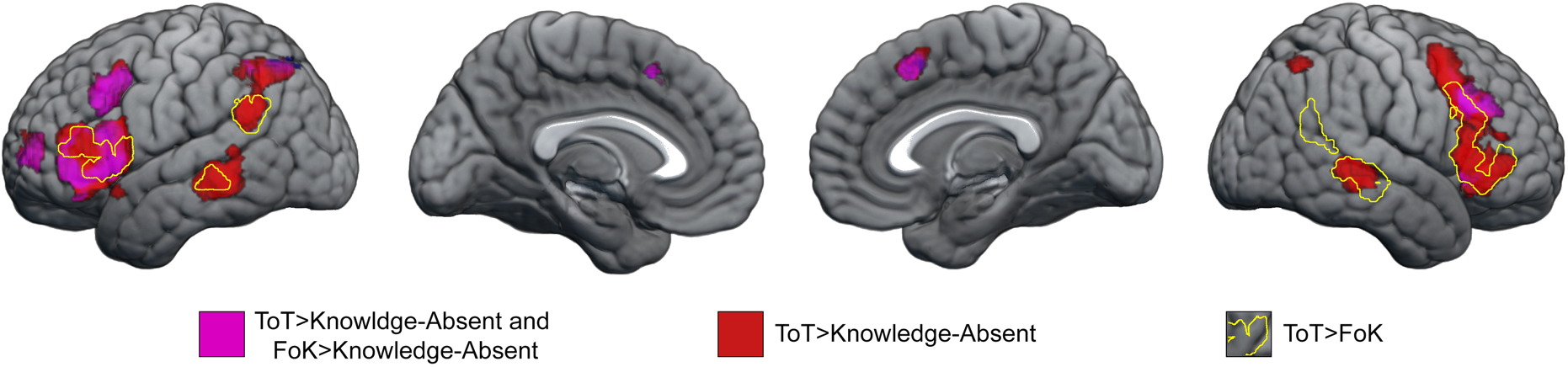
Whole-brain analysis of metacogntivie judgment comparing ToT, FoK and knoweldge-absent responses. Brain maps are initially thresholded at a voxel-wise threshold of p<.001 and cluster-level corrected p_FWE_<.05. See table 3 for significance, extent and location of clusters.

### Influence of the inclusive mask on reported results

The inclusive mask was included to clarfy the exposition of these results and focus analyses to regions previously implicated in semantic processes and did not impact the statistical signifcance of any reported results. While exluded regions were predominanrly centred in reward regions (caudate, insula, substantia nigra), the mask did diminsih the extent of some clusters. In particular, when unsuccessful access was compared to knowledge absent trials, the DMPFC cluster was more extensive and the RSC/PPA cluster extended posteriorly into the lingual gryri and cuneus. Comparing successful to unsuccessful access, additional clusters were evident in sensorimotor cortices, the postier planum temporale and cerebllum. These clusters were assoicated with a deactivation compared to fixation baseline and may result from faster reaction times for successful than unscusseful access judgments. Notably, these clustes did not persist when successful access > knowedge-absent trials (which were matched for RT) were included as a conjunction. Locations, signifcance and extent of all exluded clusters are presented here: https://figshare.com/s/194aa97ca67800f4c658.

**Table 3.** Significance, extent and location of regions during the recall event as shown in figure 6. For large clusters spanning multiple regions, local maxima are listed separately.

|  | ToT > Knowledge-Absent |  |  |  | FoK > Knowledge-Absent |  |  |  | ToT > FoK |  |  |  |
| --- | --- | --- | --- | --- | --- | --- | --- | --- | --- | --- | --- | --- |
| | cluster<br>$p(FWE-corr)$ | voxels | peak<br>T | $x,y,z$ {mm} | cluster<br>$p(FWE-corr)$ | voxels | peak<br>T | $x,y,z$ {mm} | cluster<br>$p(FWE-corr)$ | voxels | peak<br>T | $x,y,z$ {mm} |
| IFG | <.001 | 2027 | 8.69 | -50 18 0 | <.001 | 1099 | 5.07 | -52 16 4 | <.001 | 530 | 4.58 | -50 18 0 |
| ant. IFG |  |  | 7.2 | -44 42 -4 |  |  | 5.84 | -44 42 0 |  |  |  |  |
| dmPFC-SMA | .001 | 191 | 6.8 | 2 26 46 | .003 | 165 | 5.93 | -4 18 50 |  |  |  |  |
| ant. PFC | .017 | 120 | 7.05 | -40 52 4 | .027 | 109 | 5.09 | -32 60 6 |  |  |  |  |
| left premotor | <.001 | 610 | 6.9 | -42 10 38 | <.001 | 469 | 5.69 | -38 8 42 |  |  |  |  |
| right MFG | <.001 | 2193 | 6.73 | 46 22 32 | <.001 | 288 | 4.93 | 44 40 28 |  |  |  |  |
| right IFG |  |  | 6.41 | 40 24 -8 | .022 | 114 | 5.03 | 40 24 -8 | <.001 | 586 | 4.49 | 42 40 2 |
| Left IPS | <.001 | 1297 | 6.68 | -48 -48 48 | <.001 | 784 | 5.87 | -26 -56 38 |  |  |  |  |
| right mid MTG | <.001 | 628 | 6.75 | 46 -30 -2 |  |  |  |  | <.001 | 443 | 5.81 | 54 -28 -2 |
| left mid MTG | <.001 | 855 | 6.13 | -60 -36 -6 |  |  |  |  | .003 | 163 | 4.27 | -58 -34 -6 |
| right IPS | .014 | 125 | 4 | 44 -60 52 |  |  |  |  |  |  |  |  |
| cerebellum | .002 | 173 | 6.35 | -32 -62 -32 |  |  |  |  |  |  |  |  |
| cerebellum | .044 | 97 | 6.82 | -8 -78 -30 |  |  |  |  |  |  |  |  |
| left TPJ |  |  |  |  |  |  |  |  | .004 | 161 | 4.18 | -54 -44 32 |
| right TPJ |  |  |  |  |  |  |  |  | .036 | 102 | 4.43 | 60 -54 20 |

Abbreviations. IFG: inferior frontal gyrus; dmPFC-SMA: dorsomedial PFC - supplementary motor area; MFG: middle frontal gyrus; IPS: intraparietal sulcus; MTG: middle temporal gyrus; TPJ; temporoparietal junction; ant: anterior.

## Discussion

In this work, we examined the neural processes underlying successful and unsuccessful access to stored semantic knowledge. Using this paradigm, we were able to identify neural processes associated with the search for possessed knowledge and those involved in access to the relevant piece of factual information. The internal search for factual knowledge was associated with activation of both prefrontal circuitry associated with semantic control and the recruitment of cortical regions traditionally associated with place-related processing. Successful access was associated with increased activation in representational subcomponents of the semantic system.

### Recruitment of semantic control systems during successful and unsuccessful semantic access

Relative to knowledge-absent trials, unsuccessful attempts at semantic access resulted in increased activation in regions consistently implicated in semantic control: opercular and triangular sections of the IFG, and the dmPFC-SMA, consistent with a cognitive process where attempts are being made to guide selection towards the elusive semantic representation or lexical entry. While whole brain analysis showed minimal difference between successful and unsuccessful access, ROI analysis indicated increased recruitment of the pars orbitalis during successful semantic access and the dmPFC-SMA during blocked access. A more general role for the dmPFC-SMA in cognitive control may partially account for the observed dissociation. The dmPFC-SMA ROI falls within the multiple-demand network, while the left IFG ROIs do not (Fedorenko et al. 2013) and dmPFC-SMA shows parametric modulation by both working memory and semantic control demands, while left IFG exhibits modulation only by semantic control demands (Gao et al. 2021). These suggest a relatively more diverse role in cognitive control in dmPFC-SMA, that may be involved in additional extra-semantic working memory and control processes associated with blocked semantic access (Maril et al. 2005). On the other hand, the orbitalis may be more closely linked to semantic, and potentially lexical, processes, possibly though a perisylvian language-supported semantic sub-system (Xu et al. 2016, 2017).

Further recruitment of control systems in the subsequent metacognitive judgement was observed in the present study. ToTs and FoK judgment activated prefrontal systems more than knowledge-absent judgements. As in previous work, this occurred in a graded manner, with ToTs producing greater responses than FoKs (Maril et al. 2005). These effects extended beyond classically semantic prefrontal regions to the right IFG, MFG and anterior PFC, consistent with previous studies and potentially reflecting additional prefrontal cognitive processes, such as response selection or conflict resolution (Maril et al. 2001, 2005).

### Increased recruitment of representational semantic regions in successful semantic access

Compared to unsuccessful semantic access, successful access resulted in the increased recruitment in cortical regions previously associated with domain-general semantic representation, specifically, bilateral mid-MTG, the left angular gyrus, vmPFC (Binder et al. 2009; Lambon-Ralph et al. 2017). Encompassing these regions and extending over additional temporal, parietal and prefrontal areas, we observed robust selectivity for the four utilised knowledge domains, to a much greater extent than is seen with single word stimuli (Fairhall & Caramazza, 2013a). This pronounced sensitivity of sentence-level stimuli to knowledge-domain is consistent with effects seen in previous work (Rabini et al, 2021) but notably extended to show a person-selective response in a section of the right lateral fusiform gyrus anatomically overlapping the fusiform face area, a region that seldom shows person-selectivity for person-related word stimuli (Bi et al. 2016), and an object-selective response in perirhinal cortex, a region showing multivariate sensitivity to the semantic relationship between objects (Bruffaerts et al. 2013; Liuzzi et al. 2015). The scholastic knowledge-domain selectively activated core regions of the semantic system either implicated in control (left IFG) or in semantic representation (bilateral mid-MTG, Lambon-Ralph et al. 2017). Selective recruitment of left IFG is certainly consistent with scholastic information requiring additional control. However, sections of the left and right IFG are also selective for object and person knowledge-domains and the tiled nature of this selectivity across semantic domains makes it unlikely that this reflects overall differences in demands for semantic control. This pattern may indicate partial representation of semantic content within these regions or that semantic control systems are anatomically organised into subsystems that interact with different domain-selective semantic representations.

The influence of semantic access within these domain-selective regions is complex but informative. In a region-of-interest analysis, we compared the fMRI response averaged across key elements of networks exhibiting a selective response for person, place, object, and scholastic knowledge-domains. While all networks showed overall increases in activity during successful compared to unsuccessful semantic access, in networks selective for person, object and scholastic knowledge-domains, this effect was present for both preferred and non-preferred domains and was enhanced for the non-preferred domain. Notably, this pattern persists in networks that are inactive for the non-preferred categories relative to baseline during unsuccessful-access or knowledge-absent trials (i.e. person and object networks, figure 5C). Thus, while it may be intuitive to expect increases associated with semantic access to be domain-selective in domain-selective networks, the data indicates that successful access is associated with a relative broadening of the accessed semantic representations.

The place-selective network represented an exception to this pattern. In addition to showing a selective increase in response for place stimuli, these regions showed greater responses for unsuccessful-access trials than knowledge-absent trials, for both preferred and non-preferred domains. This indicates that, unlike the other knowledge-domain selective regions, PPA, TOS and RSC are involved in searching for the knowledge. These regions are implicated in knowledge-based navigation, such as locating a particular street within a university campus (Epstein et al. 2007) or locating a particular city, the provenance of a famous person or even traditional food-dish within a country (Fairhall 2020) and in making semantic judgements about kinds of people (e.g. professions, Fairhall and Caramazza 2013b). The present result extends the particular profile of place-selective semantic regions to show that they are also involved in the search for semantic information, both when access is successful and when the searched-for-fact is elusive. The reason for the particular involvement of these scene-and spatially-selective regions in the search for knowledge is uncertain. An increasing body of evidence has emerged to suggest that neural mechanism for spatial navigation in the entorhinal cortex (‘grid cells’) are repurposed to allow navigation across non-spatial dimensions, such as olfactory or pitch spaces (Aronov et al. 2017; Bao et al. 2019) and extending to navigation across certain conceptual spaces (Constantinescu et al. 2016; see Bottini and Doeller 2020 for a review; Viganò and Piazza 2020). One intriguing possibility is that the involvement of place-selective regions in navigation to complex factual information is derived via the interdependence between these regions and entorhinal cortices in spatial processing. However, this is an unexpected and preliminary finding and future work is needed to determine whether these results reflect navigation across conceptual spaces.

To facilitate clear judgements and to elicit ToT states, as in similar experimental paradigms (Maril et al. 2001, 2005), the information sought by probe questions was frequently a name or a word (e.g. ‘What is a triangle with two equal sides called?’). In this way, lexical access may contribute to the observed cortical activations, both when the answer is successfully accessed, and in unsuccessful-access trials in the form of partial lexical information (Brown and McNeill 1966; Brown 1991; Caramazza and Miozzo 1997). Lexical and semantic processes are often entangled, as access to the lexical entry of a read word is associated with access to word meaning and internal access to a lexical entry often stems from access to semantic representations. The paradigm most relevant to lexical-access processes occurring in the present study may be that of the covert/overt retrieval of lexical entries from memory. Verbal fluency paradigms (retrieving words starting with a specific letter or from a specific semantic category) predominantly recruit IFG and dmPFC (Wagner et al. 2014), while picture-cued lexical access may additionally recruit posterior aspects of the inferior/middle temporal gyri and superior temporal sulcus when lexical access is more challenging (Price 2012). Increased responses in these regions may reflect lexical (as well as semantic) processes. On the other hand, increased responses in place selective PPA, RSC and TOS, and other, particularly right hemispheric, knowledge-domain selective regions, are less likely to reflect lexical processes. However, these patterns are only suggestive. Future studies are needed to better disentangle the contribution of lexical processes to the current results.

In this work, we sought to determine the processes that distinguish successful and unsuccessful semantic retrieval. By proxy, this approach allowed us to dissociate the neural processes associated with the search for a piece knowledge from access to that knowledge. We observed that prefrontal semantic control regions are involved in the search for information and are supported by cortical regions involved in scene and spatial knowledge (PPA, TOS, and RSC) for all stimulus-domains. We saw that access to knowledge recruits representational systems, and does so broadly, recruiting both regions selective for the relevant knowledge-domain and those selective for other domains. Collectively, these results suggest that prefrontal semantic control systems and classical spatial-knowledge-selective regions work to locate relevant information and that access to a fact brings with it a broad range of associated memories incorporating other knowledge domains, rather than narrowing to information most relevant to the specific knowledge domain.

## Acknowledgements

This project was funded by the European Research Council (ERC) Starting Grant CRASK -Cortical Representation of Abstract Semantic Knowledge, awarded to Scott Fairhall under the European Union’s Horizon 2020 research and innovation program (grant agreement no. 640594).

## Notes

### Competing Interest Statement

The authors have declared no competing interest.

